# Transcriptome-wide association study of attention deficit hyperactivity disorder identifies associated genes and phenotypes

**DOI:** 10.1101/642231

**Authors:** Calwing Liao, Alexandre D. Laporte, Dan Spiegelman, Fulya Akçimen, Ridha Joober, Patrick A. Dion, Guy A. Rouleau

**Author notes:** Correspondence: Dr. Guy A. Rouleau, Montreal Neurological Institute and Hospital, Department of Neurology and Neurosurgery, 3801 University Street, Montreal, QC, Canada H3A 2B4.

## Abstract

Attention deficit/hyperactivity disorder (ADHD) is one of the most common neurodevelopmental psychiatric disorders. Previous studies have shown that the disorder is highly heritable and associated with several different risk-taking behaviors. Additionally, brain-imaging studies have identified various brain regions such as the cerebellum and frontal cortex to be altered in ADHD. Large genome-wide association studies (GWAS) have identified several loci associated with ADHD. However, understanding the biological relevance of these genetic loci has proven to be difficult. Here, we conducted the largest ADHD transcriptome-wide association study (TWAS) to date consisting of 19,099 cases and 34,194 controls and identified 9 transcriptome-wide significant hits. We successfully demonstrate that several previous GWAS hits can be largely explained by expression. Probabilistic causal fine-mapping of TWAS signals prioritized *KAT2B* with a posterior probability of 0.467 in the dorsolateral prefrontal cortex and *TMEM161B* with a posterior probability of 0.838 in the amygdala. Furthermore, pathway enrichments identified dopaminergic and norepinephrine pathways, which are highly relevant for ADHD. Finally, we used the top eQTLs associated with the TWAS genes to identify phenotypes relevant to ADHD and found an inverse genetic correlation with educational attainment and a positive correlation with “ever smoker”, maternal smoking at birth, BMI, and schizophrenia. Overall, our findings highlight the power of TWAS to identify novel risk loci and prioritize putatively causal genes.

## Introduction

Attention deficit/hyperactivity disorder (ADHD) is a common neurodevelopmental disorder globally affecting 2.5% of adults and 5% of children^1^. The disorder has been shown to be highly heritable and increases risk of substance abuse, suicide, and risk-taking behaviour^2^. Brain-imaging studies have identified various different regions, such as the cerebellum and frontal cortex, to be implicated in ADHD^3,4^. Twin studies have estimated the narrow-sense heritability of ADHD to be approximately 70%, suggesting a strong genetic component is driving the phenotypic variance^5^. Recently, a large-scale genome-wide association study (GWAS) identified several loci that were significantly associated with ADHD^6^. Despite the significant success of GWAS in delineating elements that contribute to the genetic architecture of psychiatric disorders, the loci identified are frequently difficult to characterize biologically. Often, these studies associate loci with the nearest gene, which inevitably leads to a bias for longer genes, and may not necessarily accurately depict the locus’s real effect. In contrast, transcriptomic studies have allowed for more interpretable biologically-relevant results due their use of disease-relevant cell-types and tissue, as well as the availability of databases detailing the tissue-specific expression^7^. It is also important to denote that transcriptomic studies conducted for brain disorders tend to have small sample size, by comparison to the studies of conditions where disease relevant tissue is more easily obtainable than brain tissue.

Recently, transcriptomic imputation (TI) was developed and is a powerful method to integrate genotype and expression data from large consortia such as the Genotype-Tissue Expression (GTEx) through a machine-learning approach^7^. This method derives the relationship between genotypes and gene expression to create reference panels consisting of predictive models applicable to larger independent datasets^8^. Ultimately, TI provides the opportunity to increase the statistical power to detect putative genes with small effect sizes that are associated with a disease.

To identify genetically regulated genes associated with ADHD, we used the largest ADHD cohort currently available to conduct a transcriptome-wide association study (TWAS); the cohort consists of 19,099 ADHD cases and 34,191 controls from Europe. Brain-tissue derived TI panels were used, including the 11 brain-relevant tissue panels from GTEx 53 v7 and the CommonMind Consortium (CMC). We identified 9 genes reaching within tissue panel Bonferroni-corrected significance. We additionally identify 3 novel loci and genes that were not previously implicated with ADHD. Through conditional analyses, we demonstrate that several of the genome-wide significant signals from the ADHD GWAS are driven by genetically-regulated expression. Gene set analyses of the Bonferroni-corrected TWAS genes have identified relevant pathways, among which dopaminergic neuron differentiation and norepinephrine neurotransmitter release cycle. Additionally, we queried the top eQTLs identified by TWAS in phenome databases and identified several phenotypes previously associated with ADHD such as educational attainment, body mass index (BMI), and maternal smoking around birth. Finally, genetic correlation of the pheWAS traits identified several Bonferroni-corrected significant correlations with risk-related behaviours such increased number of sexual partners and ever-smoking. In conclusion, TWAS is a powerful method that increases statistical power to identify small-effect size in genes associated with complex diseases such as ADHD.

## Results

### Transcriptome-wide significant hits

To identify genes associated with ADHD, a TWAS was conducted using FUSION^9^. A total of seventeen genes were found to be significantly associated with ADHD (Table 1, Figure 1). Here, we also detected genes that reached a within-tissue significance threshold (Supplementary Table 1). To assess inflation of imputed association statistics under the null of no GWAS association, the QTL weights were permuted to empirically determine an association statistic. The majority of genes were still significant after permutation, suggesting their signal is genuine and not due to chance.

**Table 1.**
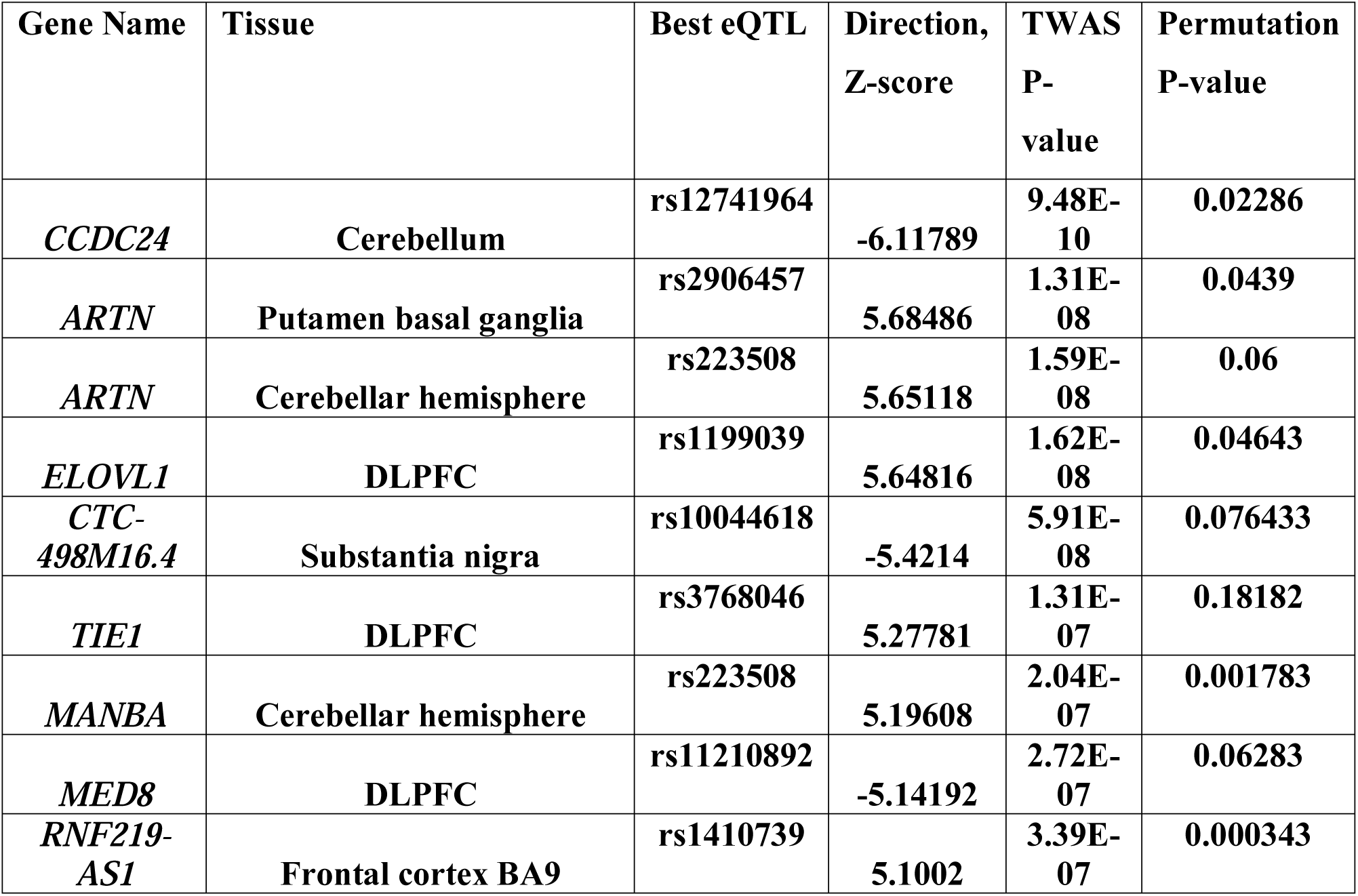
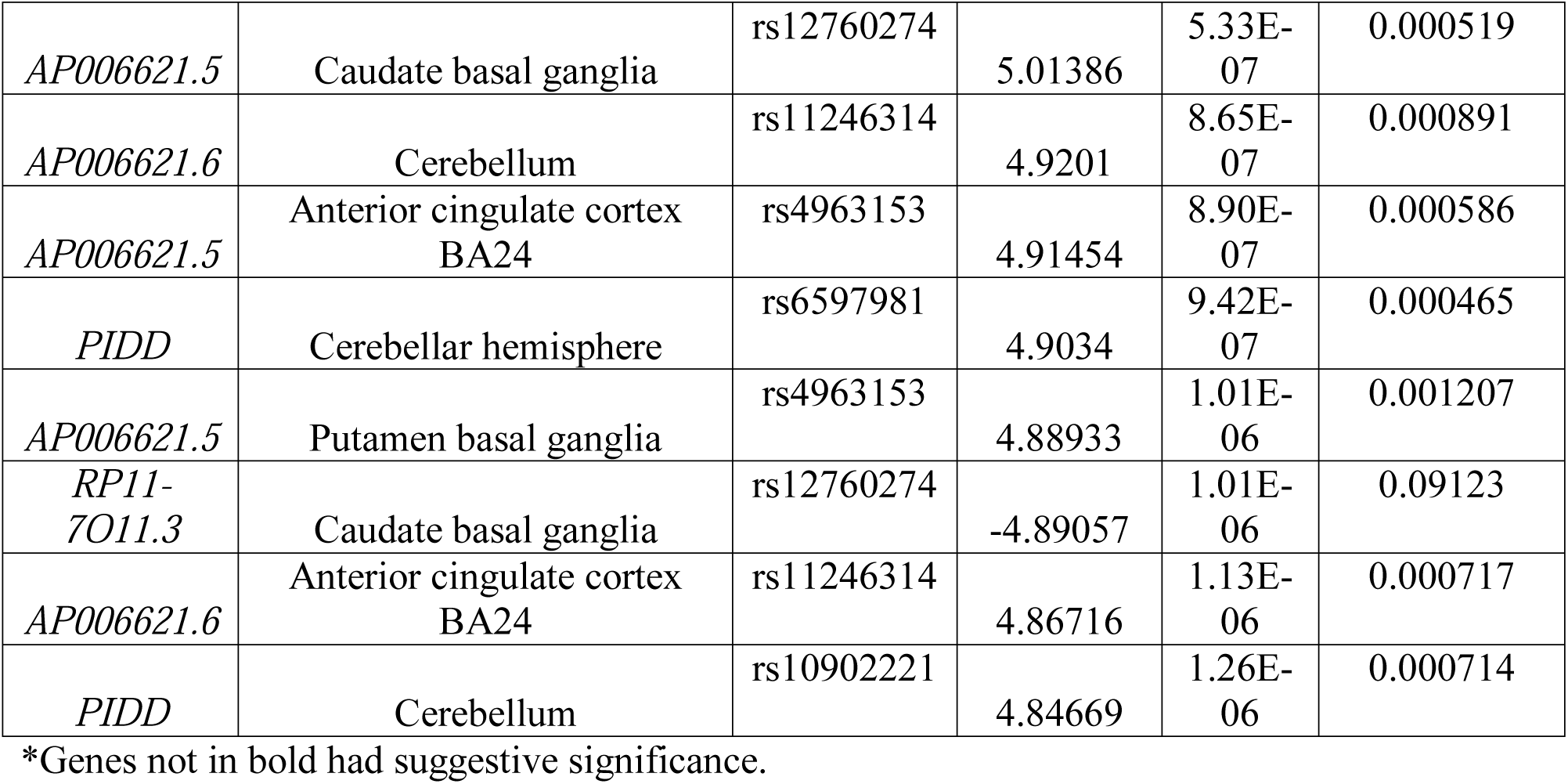
Significant TWAS genes for ADHD.

**Figure 1.**
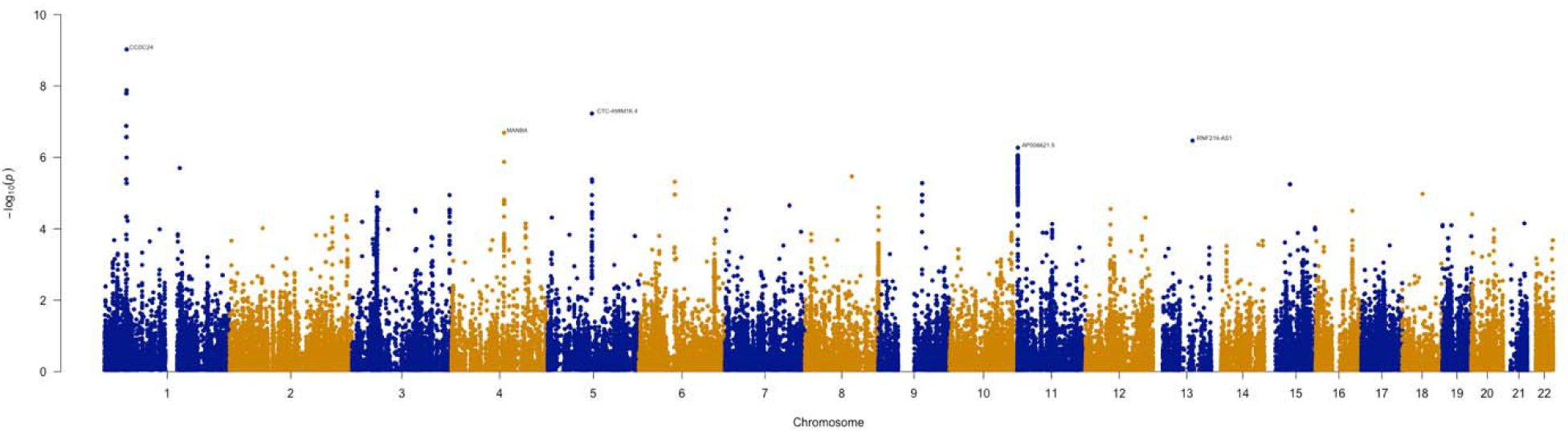
Manhattan plot of the transcriptome-wide association study for ADHD (N=19,099 cases and N=34,194 controls). Bonferroni-corrected significant genes are labelled. A significance threshold of P=4.97E-07 was used.

### Novel ADHD TWAS loci are driven completely by expression signals

Since several of the TWAS hits overlapped with significant ADHD loci, conditional and joint analyses were performed to establish whether these signals were due to multiple associated features or conditionally independent. It was observed that *AP006621.5* explains all of the signal at its loci (rs28633403 lead SNP_GWAS_ P = 4.5E-07, conditioned on *AP006621.5* lead SNP_GWAS_ P = 1) (Figure 2a). It was also found that *RNF219* explains most of the signal (rs1536776 Lead SNP_GWAS_ P= 5.5E-07, conditioned on *RNF219* to lead SNP_GWAS_ P= 5.1E-02) explaining 0.848 of the variance (Figure 2b). Conditioning on *MANBA* completely explained the variance of the loci on chromosome 4 (rs227369 Lead SNP_GWAS_ P=1.3E-07, lead SNP_GWAS_ P=1) (Figure 2c).

**Figure 2A.**
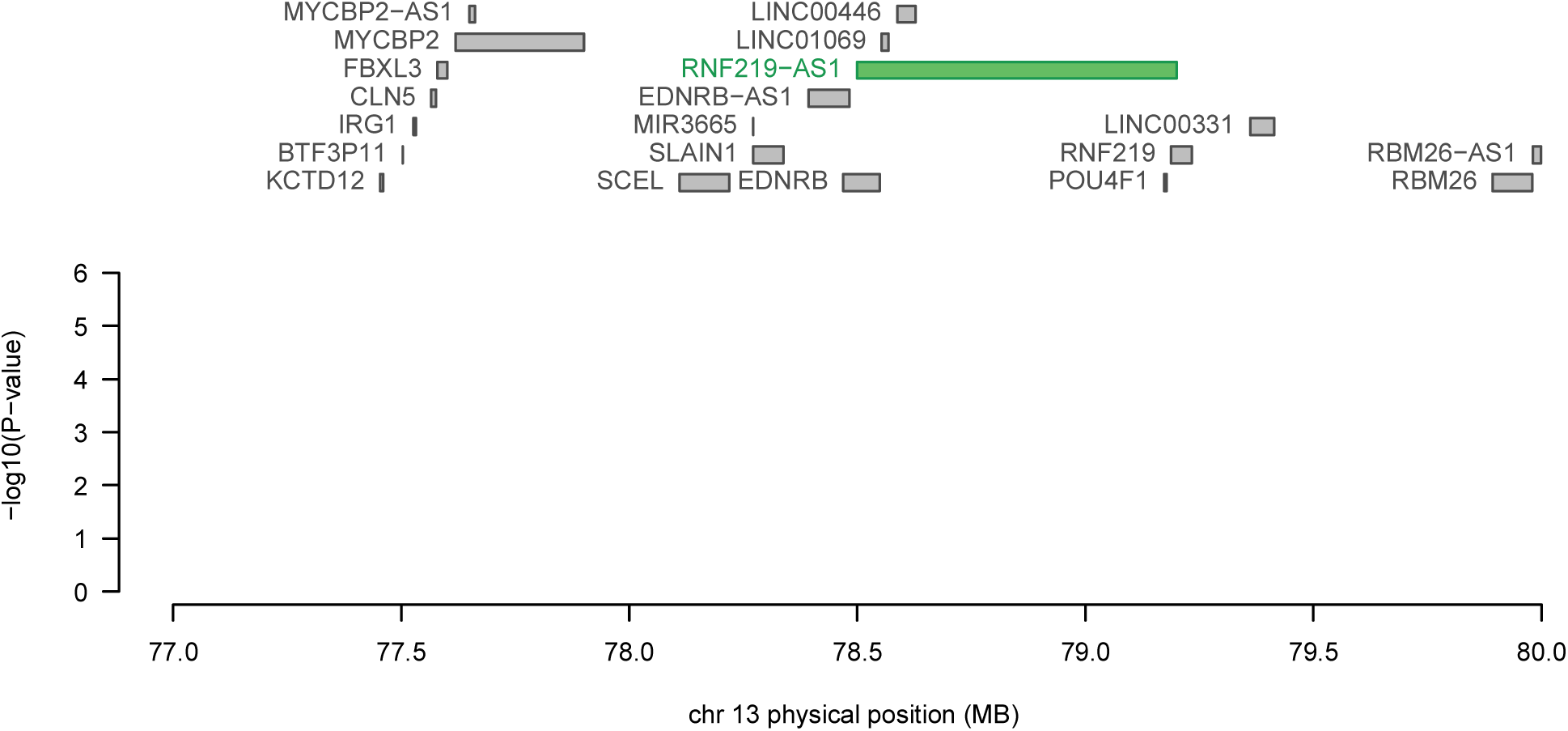
Regional association plot of chromosome 13. The top panel highlights all genes in the region. The marginally associated TWAS genes are shown in blue and the jointly significant genes are shown in green. The bottom panel shows a regional Manhattan plot of the GWAS data before (grey) and after (blue) conditioning on the predicted expression of the green genes.

**Figure 2B.**
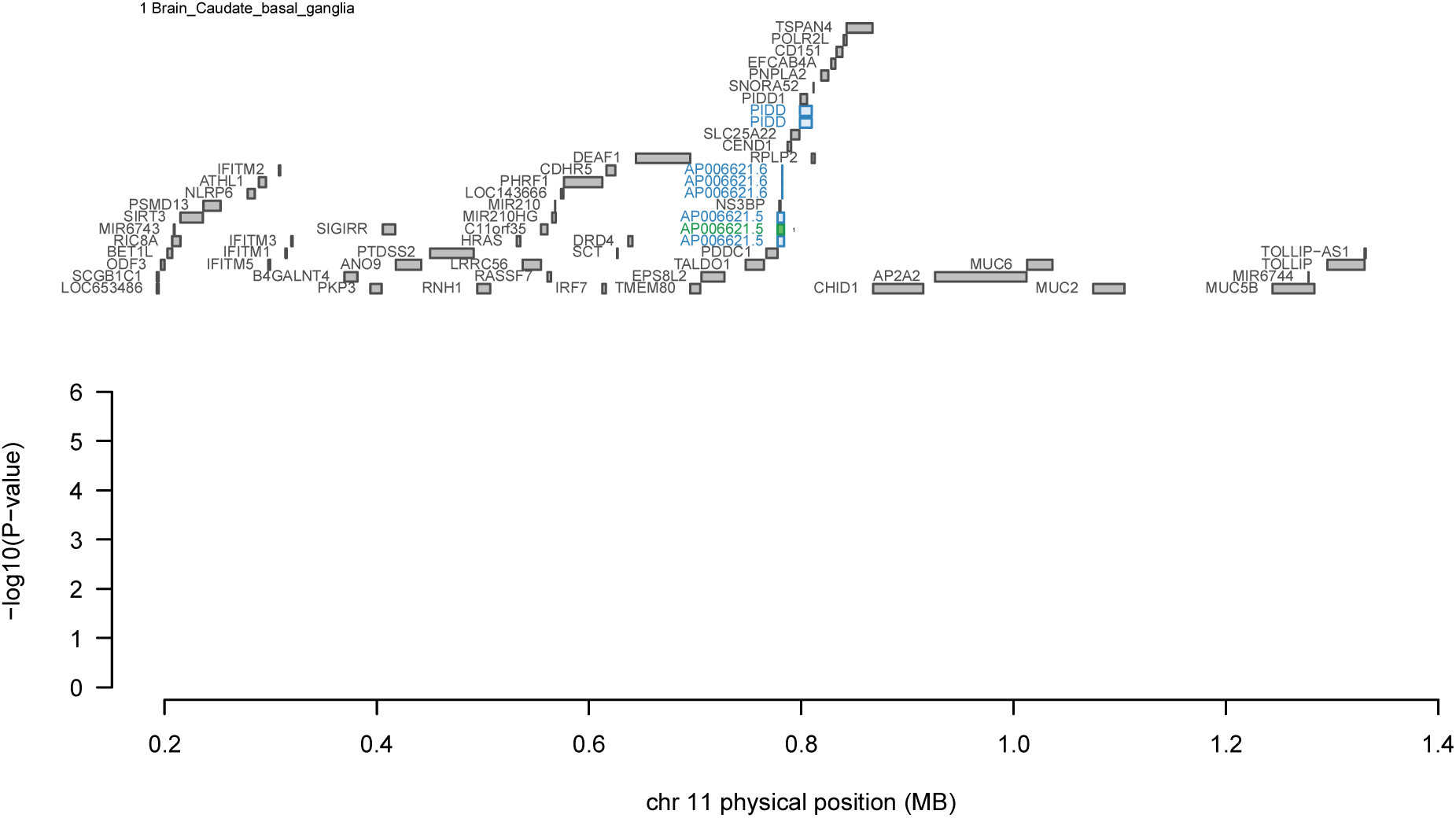
Regional association plot of chromosome 11. The top panel highlights all genes in the region. The marginally associated TWAS genes are shown in blue and the jointly significant genes are shown in green. The bottom panel shows a regional Manhattan plot of the GWAS data before (grey) and after (blue) conditioning on the predicted expression of the green genes.

**Figure 2C.**
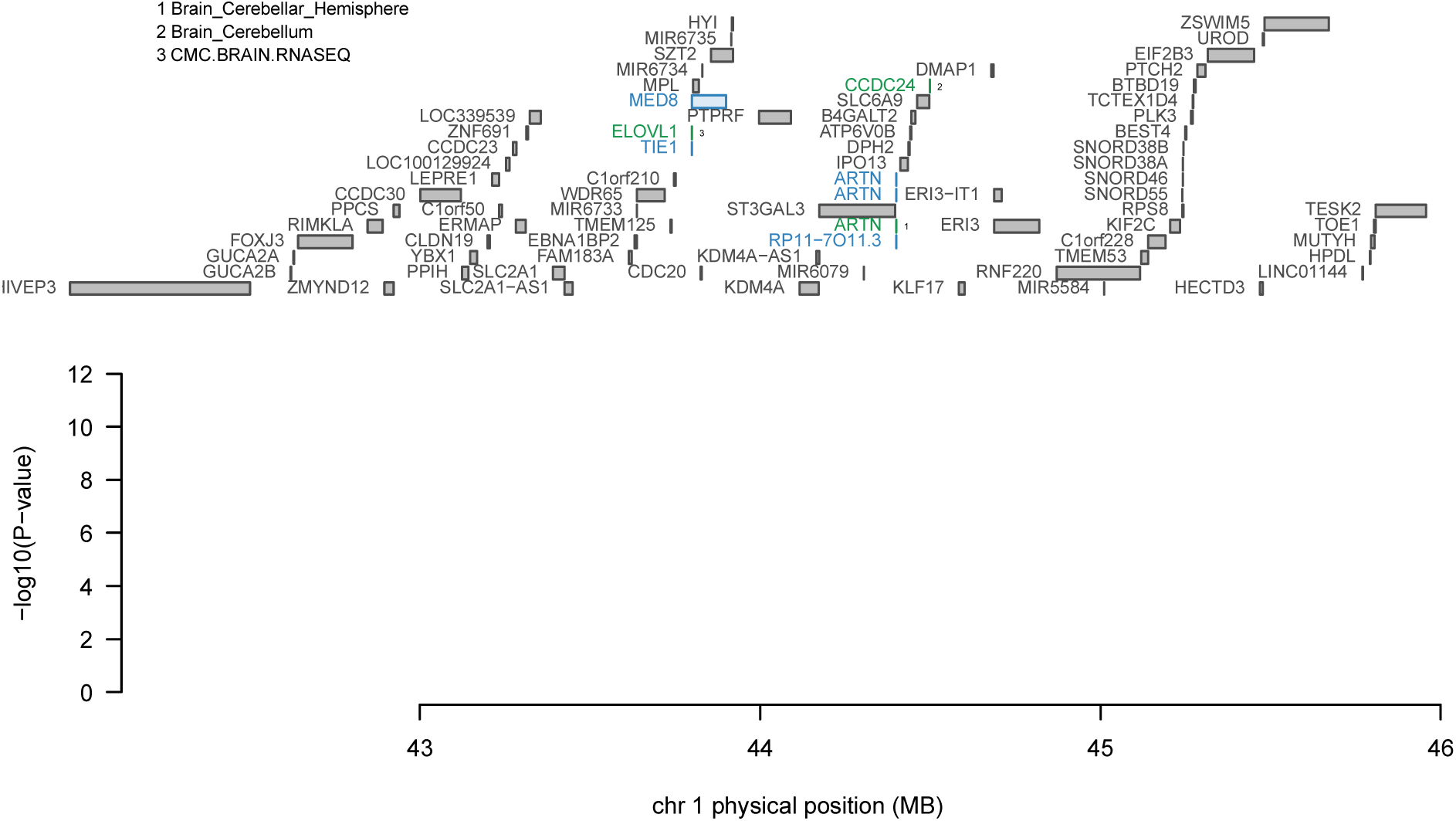
Regional association plot of chromosome 1. The top panel highlights all genes in the region. The marginally associated TWAS genes are shown in blue and the jointly significant genes are shown in green. The bottom panel shows a regional Manhattan plot of the GWAS data before (grey) and after (blue) conditioning on the predicted expression of the green genes.

**Figure 2D.**
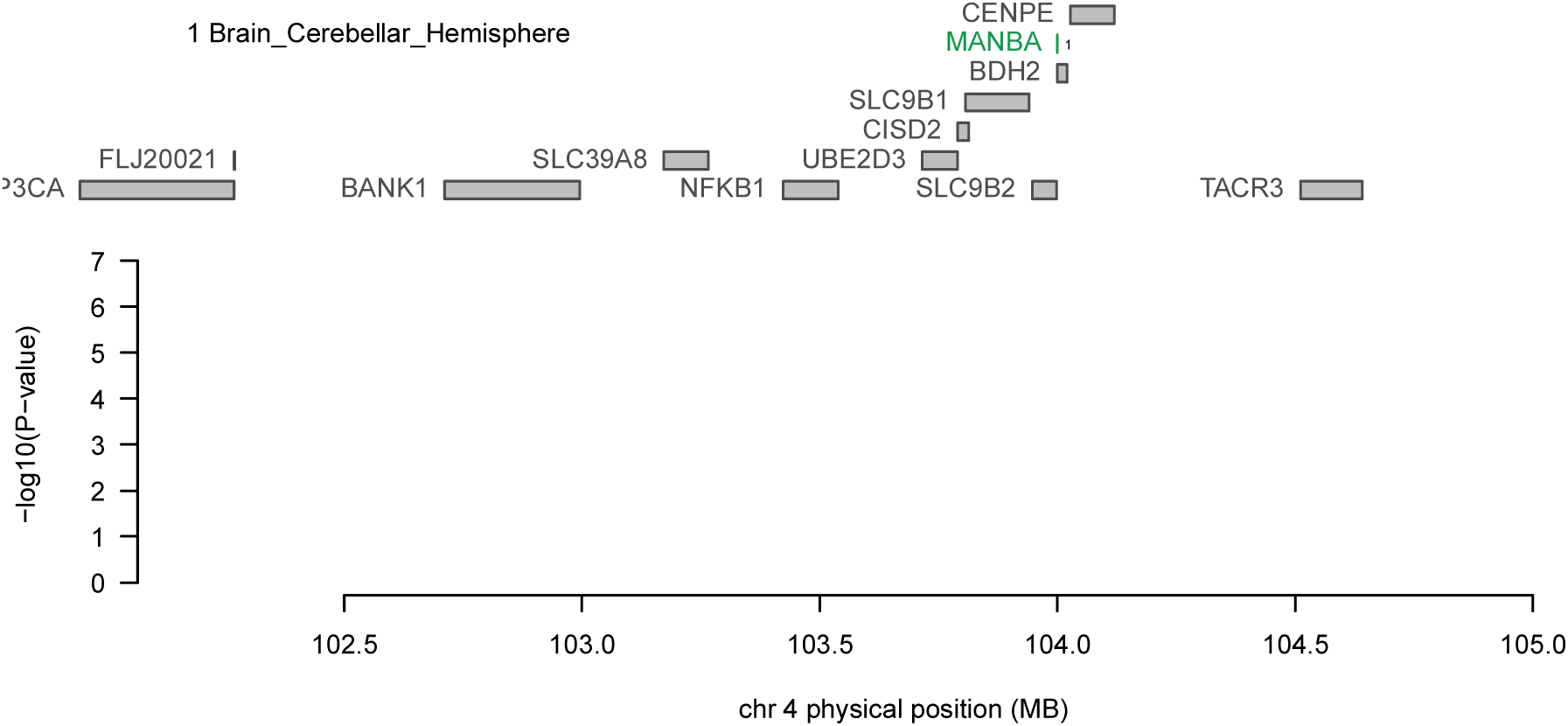
Regional association plot of chromosome 4. The top panel highlights all genes in the region. The marginally associated TWAS genes are shown in blue and the jointy significant genes are shown in green. The bottom panel shows a regional Manhattan plot of the GWAS data before (grey) and after (blue) conditioning on the predicted expression of the green genes.

**Figure 2E.**
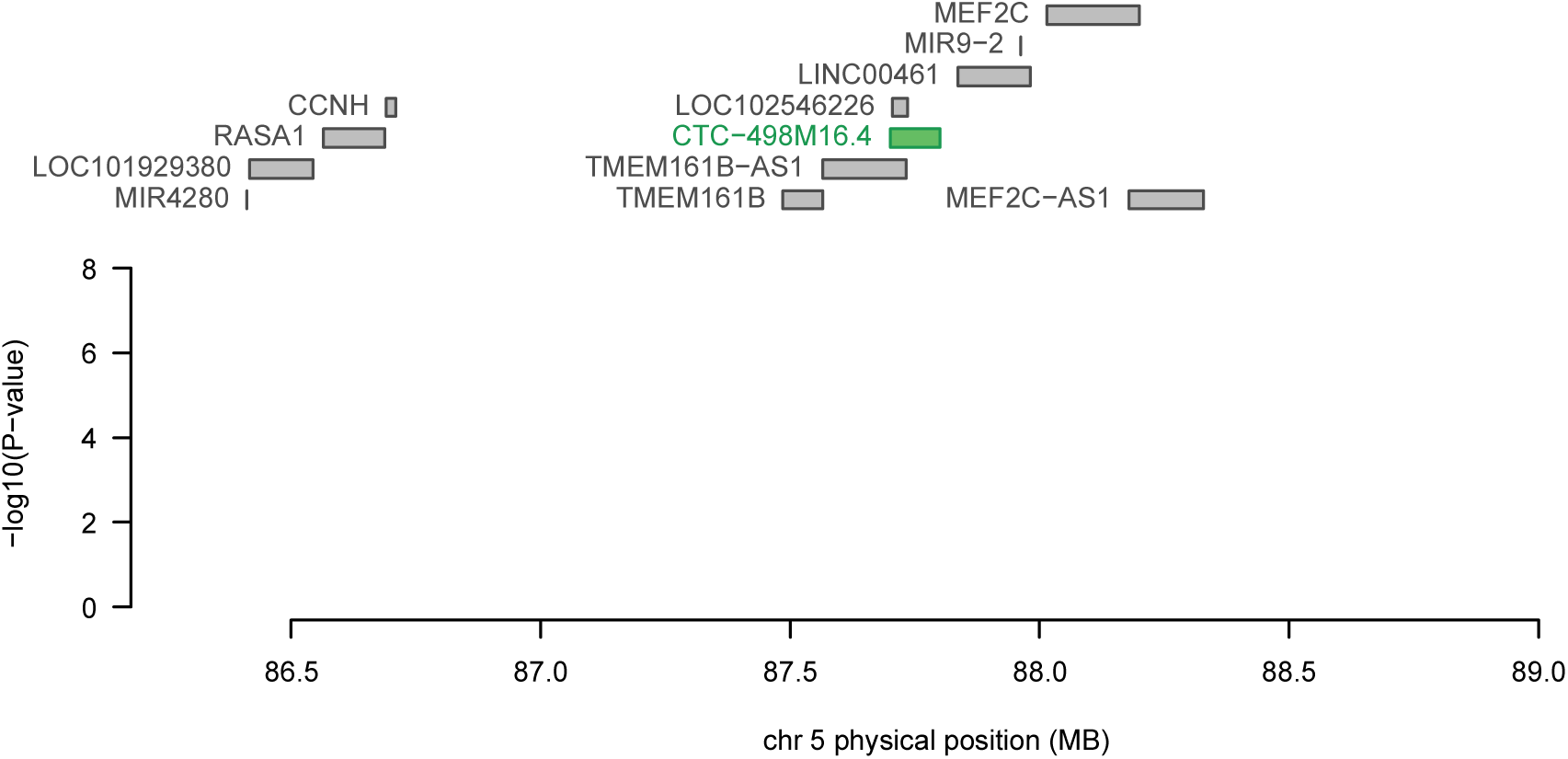
Regional association plot of chromosome 5. The top panel highlights all genes in the region. The marginally associated TWAS genes are shown in blue and the jointly significant genes are shown in green. The bottom panel shows a regional Manhattan plot of the GWAS data before (grey) and after (blue) conditioning on the predicted expression of the green genes.

### Several previously implicated ADHD loci are explained by expression signals

Similarly, conditioning on the expression of *ELOVL1, CCDC24*, and *ARTN* depending on the panel demonstrates expression-driven signals in a previously implicated ADHD loci (rs11420276 lead SNP_GWAS_ = 1.1E-12, when conditioned on *ELOVL1, CCDC24,* and *ARTN* lead SNP_GWAS_ = 7.1E-04) explaining 0.774 of the variance (Figure 2d). For the previously implicated ADHD GWAS loci at chromosome 5, conditioning on *CTC-498M16.4* explains 0.765 of the variance (rs4916723 lead SNP_GWAS_ P=1.8E-08, lead SNP_GWAS_ P=6.4E-03) (Figure 2e).

### Omnibus testing reinforces relevance of several genes

To test for whether an effect was occurring across the different panels, an omnibus test was used. There were 7 genes that passed Bonferroni-corrected significance and shown to be associated with ADHD. Interestingly, *CCDC24, ARTN, AP006621.1, CTC-498M16.4* and *MED8* remained significant. The long non-coding RNA *LINC00951* and *ST3GAL3* did not reach transcriptome-wide significance in the individual panels, but the combined omnibus score increased power to detect a signal.

### Fine-mapping of TWAS association causally implicates *TMEM161B, KAT2B* and *CTC-498M16.4*

To identify causal genes, FOCUS was used to assign a posterior inclusion probability for genes at each TWAS region and for relevant tissue types. For the genomic locus 3:20091348-3:21643707, *KAT2B* was included in the 90%-credible gene set with a posterior probability of 0.467 in the dorsolateral prefrontal cortex (Table 3). For the genomic loci 5:87390784-5:88891530, *TMEM161B, CTC-498M16.4,* and *CTC-498M16.2* were part of the credible set. The highest posterior probability for causality was 0.838 for *TMEM161B* in the amygdala and 0.139 for *CTC-498M16.4* for the hypothalamus. For the locus 16:71054116-16:72934341, *TXNL4B, HPR, DHODH, ZNF23, HP, IST1, DHX38* and *DDX19A* were included in the credible gene set. However, all the genes had lower posterior inclusion probabilities (Table 3).

**Table 2.** Omnibus significant TWAS genes for ADHD.

| Gene | Omnibus P-value |
| --- | --- |
| ***ARTN*** | **3.27E-11** |
| ***CCDC24*** | **3.90E-10** |
| ***LINC00951*** | **2.01E-08** |
| ***ST3GAL3*** | **3.27E-07** |
| ***AP006621.1*** | **1.17E-06** |
| ***MED8*** | **1.60E-06** |
| ***CTC-498M16.4*** | **2.00E-06** |
| *MST1R* | 5.99E-06 |
| *AP006621.5* | 1.12E-05 |

\*Genes not in bold had suggestive significance.

**Table 3.**
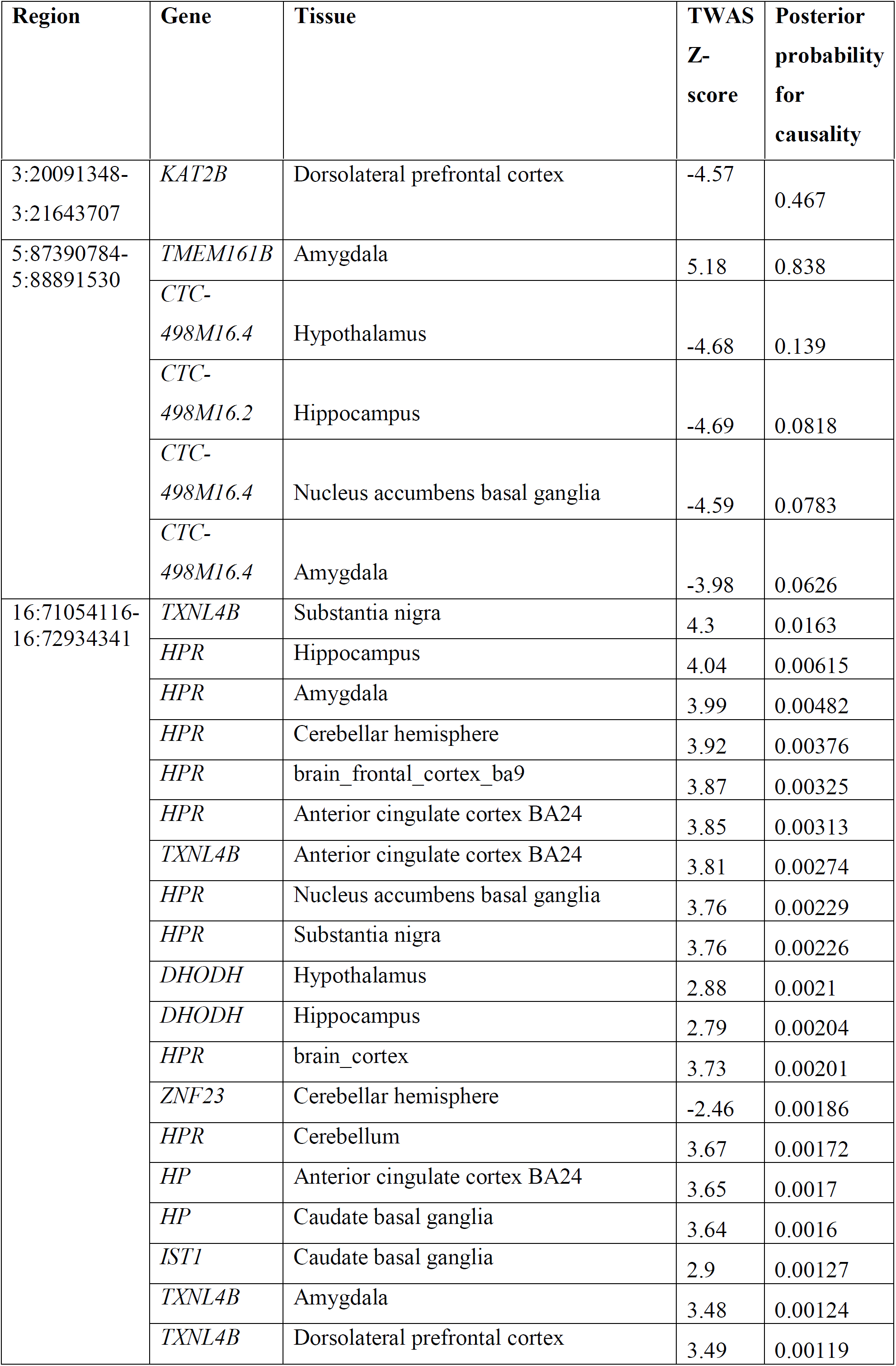
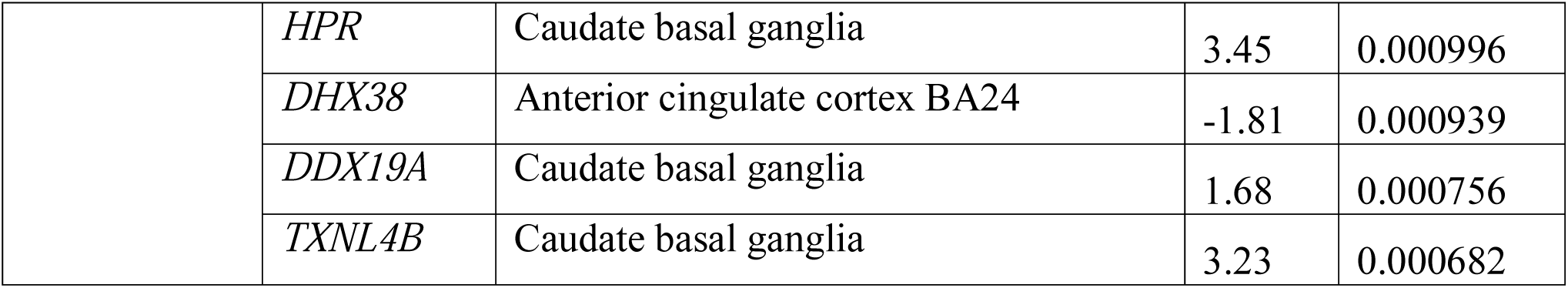
Causal posterior probabilities for genes in 90%-credible sets for available ADHD GWAS regions.

### Pathway enrichment highlights importance of lipid metabolism, dopaminergic neurons and norepinephrine

To understand the biologically relevant pathways from the transcriptome-wide significant hits, pathway and gene ontology analyses were conducted using Reactome and GO. The genes grouped into three different clusters based on co-expression of public RNA-seq data (N=31,499) (Supplemetnary Figure 2). Several relevant pathways were significantly enriched such as dopaminergic neuron differentiation (P=3.5E-03), norepinephrine neurotransmitter release cycle (P=4.4E-03), and triglyceride lipase activity (P=2.9E-03) when analyzing all genes together. Interestingly, several relevant cellular regions such as the axon and dendritic shaft were also enriched (Supplementary Table 2).

### Phenome-wide association study (pheWAS) of top eQTLs identified clinically relevant phenotypes

To understand phenotypes that may be associated or co-morbid with ADHD, a pheWAS was done for each eQTL (Supplementary Table 3). Since most eQTLs were associated with ADHD, we chose to exclude it from Table 3 to emphasize the other three top phenotypes per SNP. Several risk-associated phenotypes such as “ever-smoker”, alcohol intake over 10 years, and maternal smoking around birth were found to be significantly associated with the eQTLs.

### Genetic correlation of pheWAS traits demonstrate the relevance of metabolic and behavioural phenotypes

To determine whether the pheWAS traits were genetically correlated and in which direction, genetic correlation was done between the most recent (as of the writing of this publication) GWAS for each of the phenotypes. Interestingly, there was a strong negative correlation between educational attainment (Supplementary Figure 1). Furthermore, a positive correlation between maternal smoking around birth, body mass index, ever smoker, and schizophrenia.

## Discussion

Attention deficit/hyperactivity disorder is a common disorder that affects millions of people worldwide. While recent GWAS has been successful and identifying risk loci associated with ADHD, the functional significance of these associations continue to remain elusive due to the inability to fine-map to tissue-specific and -relevant genes. Here, we conducted the largest ADHD TWAS to date, using the summary statistics of over 50,000 individuals from the most recent ADHD GWAS. This approach creates genotype-expression reference panels using public consortia through machine learning approaches, allowing for imputation and association testing of independent large-scale data^7,8^. We identified 9 genes-associated with ADHD risk and different tissue types, localizing to 5 different regions in the genome. Interestingly, conditional and joint analyses demonstrated that the TWAS expression signals were driving the significance for several previously implicated ADHD loci when conditioned on the top TWAS gene. For smaller genes such as *AP006621.5 gene,* they would normally go unnoticed due to the many larger protein-coding genes nearby. However, our TWAS results demonstrated that the expression of *AP006621.5* fully explained the suggestive ADHD GWAS signal, highlighting the power of TWAS to fine-map towards genes of interest. Moreover, across all brain tissue types, the significant hits were consistently seen in the following biologically relevant tissue for ADHD: cerebellum, dorsolateral prefrontal cortex, frontal cortex, basal ganglia, and anterior cingulate cortex. These regions are consistent with previously implicated deficit points in the frontal-subcortical catecholamine and dopamine networks for ADHD^10,11^. However, many TWAS hits tend to be correlated due to co-expression. Causal gene prioritization programs such as using FOCUS probabilistically help fine-map towards credible genes^12^.

Fine-mapping of TWAS hits included *KAT2B* in the credible-set with a posterior probability of 0.467 in the dorsolateral prefrontal cortex. The literature shows that ADHD has been associated with weaker function of the prefrontal cortex compared to healthy individuals^13^. *KAT2B* is a lysine histone acetyltranferase highly expressed in the brain^14^. Previous evidence has suggested that lysine acetylation is importance for brain function and proper development^14^. At another locus, *TMEM161B*, a transmembrane protein, had the highest posterior inclusion probability of 0.838 in the amygdala. Brain imaging studies have shown that the amygdala has decreased volume in ADHD patients^15^. Genetic variants in the gene have also been previously associated with major depressive disorder (MDD), which is a disorder that is often co-morbid with ADHD^16^. Furthermore, genetic correlation of ADHD and MDD has been shown to have a significant positive genetic correlation^17^. Another gene at this locus, *CTC-498M16.4*, was included in the credible-set for multiple relevant brain tissue types such as the hypothalamus, hippocampus and amygdala as well. However, the posterior inclusion probability was lower for this gene.

Interestingly, pathway and GO enrichment reinforced several pathways that have previously been reported as biologically relevant. Both dopaminergic and noradrenergic contributions have been implicated in the pathogenesis of ADHD^11^. For the top eQTLs associated with each transcriptome-wide significant gene, many re-occurring phenotypes relevant to ADHD were present such as “ever-smoker” and number of sexual partners. A genetic correlation between those available traits from public GWAS data and the most recent ADHD GWAS found inverse correlation for education attainment, consistent with studies on educational outcome with ADHD. Additionally, a genetic correlation for risky behaviours such as “ever-smoker” and maternal smoking around birth were positively correlated. Maternal smoking has often been suggested to be a risk factor for ADHD^18^. However, the positive genetic correlation could suggest pleiotropy for genetic loci associated with both phenotypes. This would be consistent with pheWAS results showing that some eQTLs were highly associated with both ADHD and smoking. Future studies should investigate segregating large heterogeneous cohorts on the phenotypes associated with these eQTLs to identify subgroups of the disease.

We conclude this study with several caveats and potential follow-up studies. First, TWAS associations could potentially be due to confounding because the gene expression levels that were imputed are derived from weighted linear combinations of SNPs. These SNPs could be included in non-regulatory mechanisms driving the association and risk, ultimately inflating certain statistics. Although the permutation tests and probabilistic fine-mapping used in this study try to protect against these spurious chance events, there is still a possibility of this occurring. Second, a follow-up study will require a large replication cohort, which may be difficult to ascertain since this current largest GWAS dataset was used in this study. Future studies could investigate the possibility of using gene-risk scores in additional cohorts to validate any findings from this study. Finally, a given gene may have other regulatory features that do not go through eQTLs and still have downstream effect on the trait. Here, we successfully managed to identify several putatively causal genes such as *TMEM161B* and *KAT2B* associated with ADHD. To conclude, TWAS is a powerful statistical method to identify small and large-effect genes associated with ADHD and helps with understanding the molecular underpinning of the disease.

## Methods

### Genotype data

Summary statistics were obtained through the ADHD Workgroup of the Psychiatric Genomics Consortium (PGC-ADHD)^19^. Details pertaining to participant ascertainment and quality control were previously reported by Demontis *et al*. (2019)^6^. The data used in this paper includes only the European population from the ADHD GWAS (N=19,099 cases and N=34,194 controls).

### Transcriptomic imputation

Transcriptomic imputation (TI) was done using eQTL reference panels derived from tissue-specific gene expression coupled with genotypic data. Here, we used 10 brain tissue panels from GTEx 53 v7 and the CommonMind Consortium (CMC)^20^. A strict Bonferroni-corrected study-wise threshold was used: P=4.97E-07 (0.05/100,572) (total number of genes across panels). FUSION was used to conduct the transcriptome-wide association testing. The 1000 Genomes v3 LD panel was used for the TWAS. FUSION utilizes several penalized linear models such as GBLUP, LASSO, Elastic Net^7^. Additionally, a Bayesian sparse linear mixed model (BSLMM) is used. FUSION computes an out-sample R^2^ to determine the best model by performing a fivefold cross-validating of every model. Further details can be read about from the original manuscript. After, a multiple degree-of-freedom omnibus test was done to test for effect in multiple reference panels. This test will account for pairwise correlation between functional features. The threshold for the omnibus test was P=4.64E-06 (0.05/10,323) (number of genes tested for omnibus).

### Conditionally testing GWAS signals and permutation

To determine how much GWAS signal remains after the expression association from TWAS is removed, joint and conditional testing was done for genome-wide Bonferroni-corrected TWAS signals. The joint and conditional analyses help to determine genes with independent genetic predictors associated with ADHD from genes that are simply co-expressed with a genetic predictor. Each ADHD GWAS SNP association was conditioned on the joint gene model one SNP at a time. To assess inflation of imputed association statistics under the null of no GWAS association, a permutation test (N=100,000 permutations) was conducted to shuffle the QTL weights and empirically determine an association statistic. Permutation was done for each of the significant loci. The loci that pass the permutation test demonstrate levels of heterogeneity captured by expression and are less likely to be co-localization due to chance. It should be noted that the permutated statistic is very conservative and truly causal genes could fail to reject the null due to the QTLs having complex and high linkage disequilibrium.

### Fine-mapping of TWAS associations

To address the issue of co-regulation in TWAS, we used the program FOCUS (Fine-mapping of causal gene sets) to directly model predicted expression correlations and to give a posterior probability for causality in relevant tissue types^12^. FOCUS identifies genes for each TWAS signal to be included in a 90%-credible set while controlling for pleiotropic SNP effects. The same TWAS reference panels for FUSION were used.

### Gene-set analyses

Due to the stringent Bonferroni-corrected significance, we relaxed the threshold for pathway analyses since Bonferroni-correction assumes independence and genes tend to be correlated due to co-expression. Gene clustering was done using the GeneNetwork v2.0 RNA sequencing database (N=31,499)^21^. Genes meeting a Bonferroni significance threshold of P= 2.98E-06 (0.30/100,572) was used. Agnostic analyses of pathways in databases such as Reactome and GO were done to identify pathways relevant to ADHD.

### Phenome-wide association studies

To identify phenotypes associated with the top eQTL for each TWAS gene, a phenome-wide association study (pheWAS) was done for each SNP. The top three phenotypes (excluding ADHD) were reported. PheWAS was done using public data provided by GWASAtlas.

### Genetic correlation

To determine the genetic relationship between ADHD and the phenotypes identified from pheWAS, genetic correlation of the traits was done for available GWAS data. The most recent GWAS data (as of 2019) for each trait was used for the correlation. The significance threshold was corrected for the number of tested traits with a Bonferroni-correction.

## Acknowledgements

We would like to acknowledge Jay Ross, Cynthia Bourassa, Nargess Farhang, Elias Jabbour for their scientific advice and editing of the manuscript.

