## Supplementary Figures and Tables for "Transcriptome-wide association study of attention deficit hyperactivity disorder identifies associated genes and phenotypes"

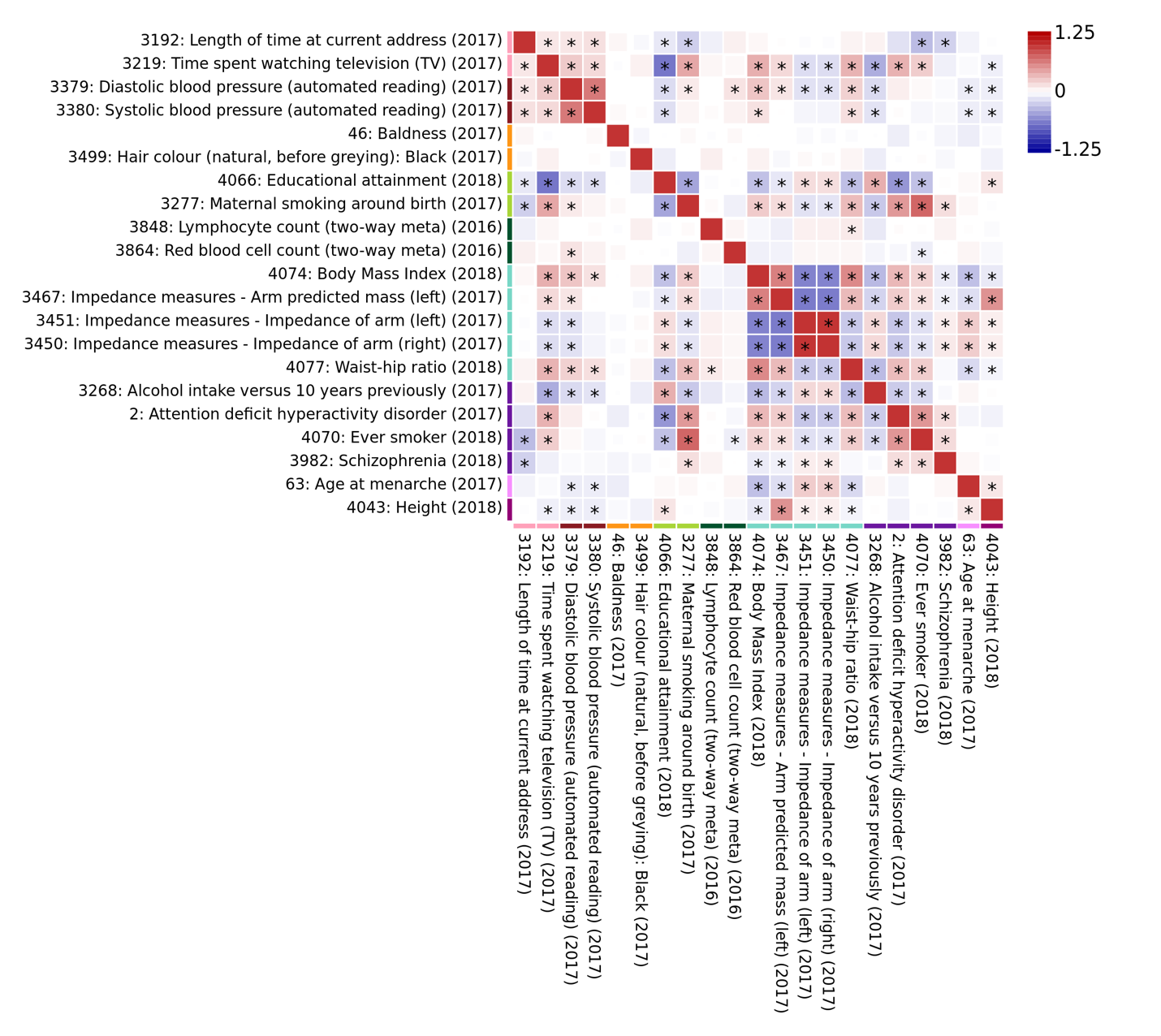

**Supplementary Figure 1. Genetic correlation plot of phenotypes associated with top ADHD eQTLs**. An asterisk in the box indicates the correlation passes Bonferroni significance threshold. Phenotypes are clustered by domain and derived from public genome-wide association study summary statistics.

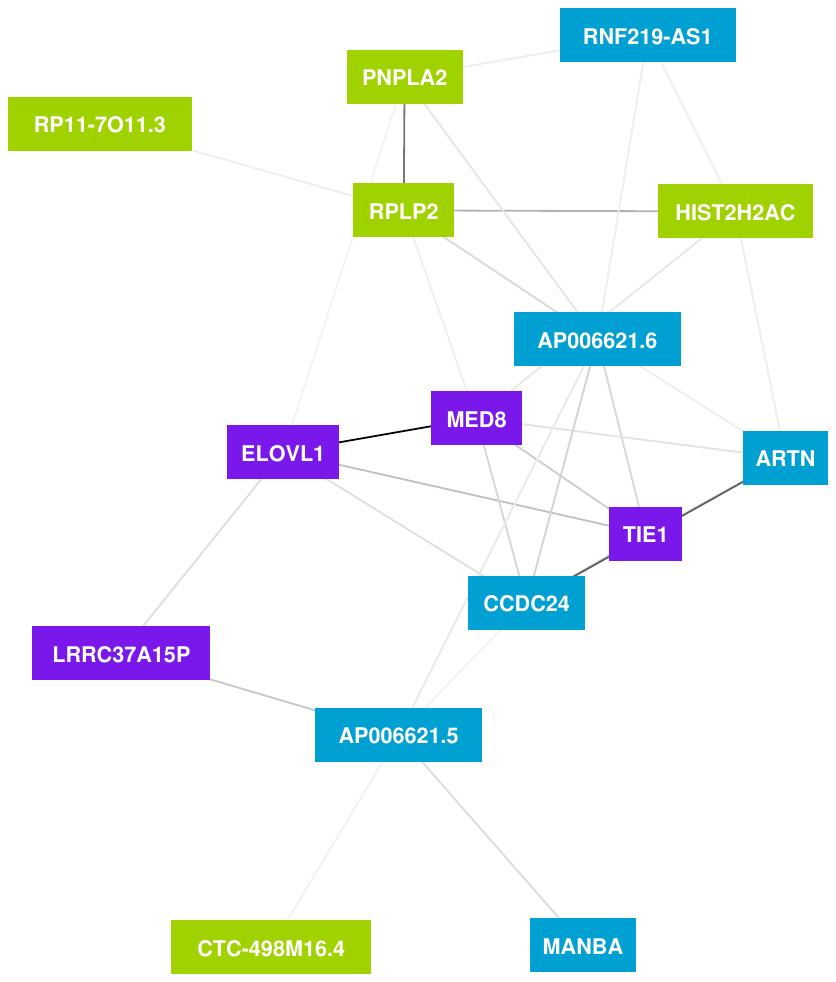

**Supplementary Figure 2. Gene clustering of differentially expressed genes for the suggestive TWAS genes based on gene co-expression.** (A) Public RNA sequencing data (N=31,499) was used to determine co-expression profiles. Gene cluster 1 identified in blue. Gene cluster 2 identified in green. Gene cluster 3 identified in purple.

**Supplementary Table 1. Within-tissue panel significance thresholds**

| **TI Panel** | **Number of genes** | **Within-tissue significance threshold** |
| --- | --- | --- |
| GTEx Brain Anterior Cingulate Cortex ba24 | 8731 | 5.73E-06 |
| GTEx Caudate Basal Ganglia | 9145 | 5.47E-06 |
| GTEx Cerebellar Hemisphere | 9451 | 5.29E-06 |
| GTEx Cerebellum | 10002 | 5.00E-06 |
| GTEx Cortex | 9162 | 5.46E-06 |
| GTEx Frontal Cortex BA9 | 9031 | 5.54E-06 |
| GTEx Hippocampus | 8535 | 5.86E-06 |
| GTEx Hypothalamus | 8551 | 5.85E-06 |
| GTEx Nucleus Accumbens Basal Ganglia | 8913 | 5.61E-06 |
| GTEx Brain Putamen Basal Ganglia | 8759 | 5.71E-06 |
| CMC DLPFC | 10292 | 4.86E-06 |
| Total | 100572 | 4.9E-07 |

**Supplementary Table 2. Significant pathways of TWAS genes identified through gene network analysis.**

| **Pathway** | **Significance** | **Database** |
| --- | --- | --- |
| Dopaminergic neuron differentiation | 3.5E-03 | GO Processes |
| Delayed rectifier potassium channel activity | 8.8E-04 | GO Function |
| Dendritic shaft | 1.7E-03 | GO Cellular |
| Axon Terminus | 3.4E-03 | Go Cellular |
| Neurotransmitter release cycle | 4.4E-03 | Reactome |
| Norepinephrine neurotransmitter release cycle | 5.5E-03 | Reactome |
| Triglyceride lipase activity | 2.9E-03 | GO Function |

**Supplementary Table 3. Phenotypes associated with top eQTLs derived from TWAS**

| **dbSNP or dbSNP ID** | **Phenotypes excluding ADHD (P-value)** |
| --- | --- |
| rs12741964 | 1. Red blood cell count (0.0001068091)  2. Educational attainment (0.000458)  3. Time spent watching TV (0.0003059) |
| rs2906457 | 1. Educational attainment (1.69E-10) 2. Alcohol intake versus 10 years previously (1.52E-8)  3. Cooked vegetable intake (8.41E-08) |
| rs223508 | 1. Lymphocyte count (2.21E-09) 2. Impedance of right arm (3.05E-09) 3. Impedance of left arm (5.89E-09) |
| rs1199039 | 1. Systolic blood pressure (9.24E-13) 2. Diastolic blood pressure (8.14E-12) 3. Height (3.6E-11) |
| rs10044618 | 1. Number of sexual partners (4.23E-12) 2. Educational attainment (6.08E-11) 3. Left arm predicted mass (1.2E-09) |
| rs3768046 | 1. Systolic blood pressure (7.61E-13) 2. Diastolic blood pressure (1.24E-11) 3. Height (9.1E-11) |
| rs223508 | 1. Lymphocyte count (2.21E-09) 2. Impedance of right arm (3.05E-09) 3. Impedance of left arm (5.89E-09) |
| rs11210892 | 1. Educational attainment (6.4E-19) 2. Age of menarche (5.47E-14) 3. Ever smoker (3.57E-13) |
| rs1410739 | 1. Hair colour (black) (0.00003779) 2. Baldness (0.000128406) 3. Hair colour (0.000128406) |
| rs12760274 | 1. Educational attainment (1.16E-09) 2. Maternal smoking around birth (4.37E-07) 3. Length of time at current address (0.000008061) |
| rs11246314 | 1. Waist-hip ratio (1.49E-13) 2. Impendance of right leg (2.35E-11) 3. Impedance of left leg (9.84E-11) |
| rs4963153 | 1. Waist-hip ratio (1.58E-13) 2. Body mass index (5.6E-09)  3. Impedance of right leg (6.64E-09) |
| rs6597981 | 1. Waist-hip ratio (8.69E-14) 2. Impedance of right leg (1.68E-10)  3. Waist-hip ratio (2.02E-10) |
| rs4963153 | 1. Waist-hip ratio (1.58E-13) 2. Body mass index (5.6E-09) 3. Impedance measures (6.64E-09) |
| rs12760274 | 1. Educational attainment (1.16E-09) 2. Maternal smoking around birth (4.37E-07) 3. Schizophrenia (0.00000905) |
| rs10902221 | 1. Waist-hip ratio (1.85E-13) 2. Impedance of right leg (3.22E-11) 3. Impedance of left leg (1.03E-10) |
